## Supplementary File1 - Supplementary Figures for "Residues Neighboring an SH3-Binding Motif Participate in the Interaction *In Vivo*"

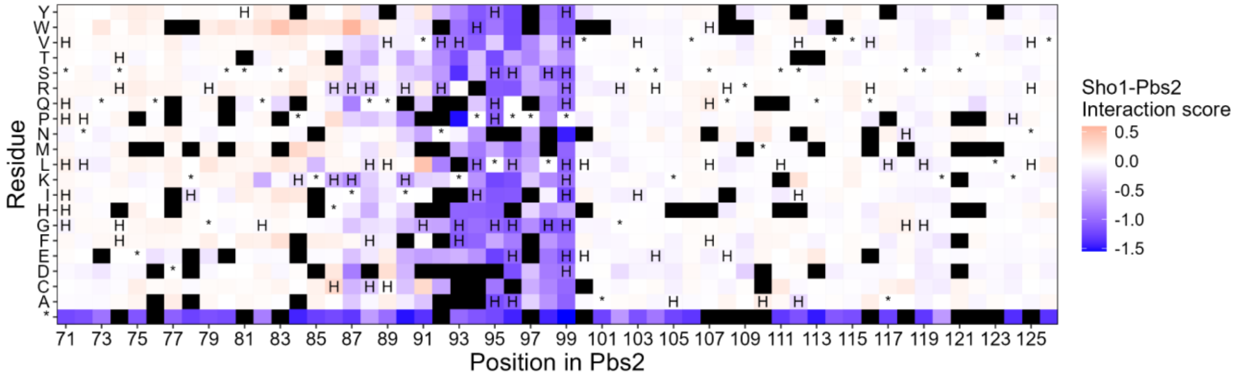


Figure S1.

Sho1-Pbs2 interaction scores for Pbs2 mutants in the DHFR-PCA on the DMS library of the region surrounding the Pbs2 motif (positions 71-126). Scores are indicated for each mutant according to their position in Pbs2 and the residue that it was mutated to. The wild-type residue for each position is indicated by an asterisk (*). Mutants which significantly affect the Hog1-Pbs2 interaction are marked with the letter H (Two-sided Mann-Whitney U test with false discovery rate corrected p-value < 0.05). Mutants which were removed from the dataset for having too few reads in the initial timepoint are marked in black. Each measurement is the median of all codons in each replicate coding for the same residue, with 3 to 18 replicates for each mutant.

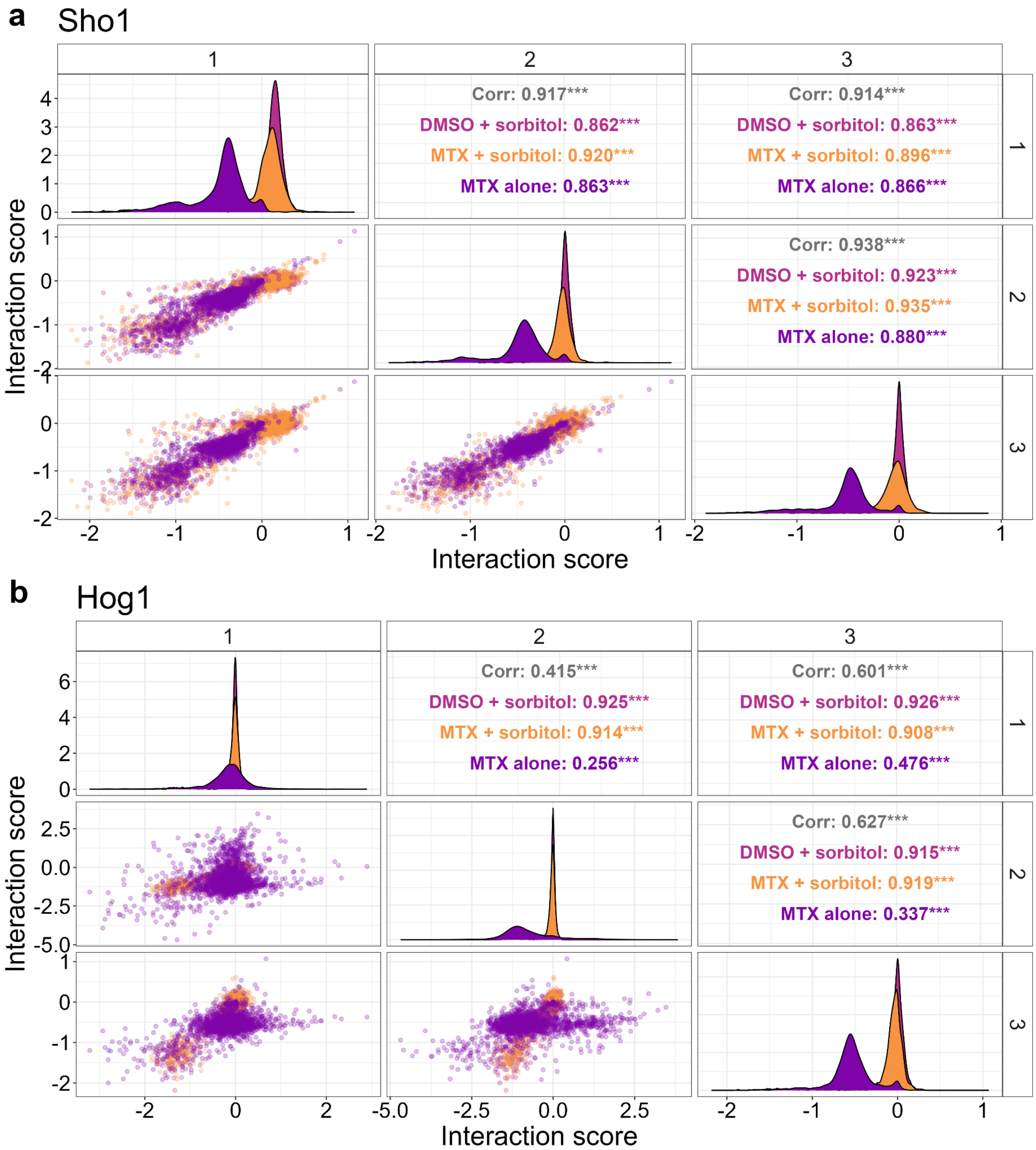

a
Sho1
Corr: 0.917***
DMSO + sorbitol: 0.862***
MTX + sorbitol: 0.920***
MTX alone: 0.863***
Corr: 0.914***
DMSO + sorbitol: 0.863***
MTX + sorbitol: 0.896***
MTX alone: 0.866***
Corr: 0.938***
DMSO + sorbitol: 0.923***
MTX + sorbitol: 0.935***
MTX alone: 0.880***
Interaction score
Interaction score
b
Hog1
Corr: 0.415***
DMSO + sorbitol: 0.925***
MTX + sorbitol: 0.914***
MTX alone: 0.256***
Corr: 0.601***
DMSO + sorbitol: 0.926***
MTX + sorbitol: 0.908***
MTX alone: 0.476***
Corr: 0.627***
DMSO + sorbitol: 0.915***
MTX + sorbitol: 0.919***
MTX alone: 0.337***
Interaction score
Interaction score

Figure S2.

Correlation between replicates in the DHFR-PCA experiments on DMS libraries surrounding region of the Pbs2 binding motif a) Scatterplot between the 3 replicates of the Sho1-Pbs2 DHFR-PCA interaction scores for mutants in the DMS library of the region surrounding the Pbs2 motif (positions 71-126). Each point represents one Pbs2 codon variant in one condition (with or without sorbitol, with MTX or with only DMSO), as indicated by the color in the corresponding panel on the top right. Pearson correlations also indicated in the top right panels. In the panels on the diagonal, the distribution of scores for a given condition and replicate is shown. The replicate number is indicated above and to the right of the plots. b) Scatterplot between the 3 replicates of the Hog1-Pbs2 DHFR-PCA interaction scores for mutants in the DMS library of the region surrounding the Pbs2 motif (positions 71-126). Each point represents one Pbs2 codon variant in one condition (with or without sorbitol, with MTX or with only DMSO), as indicated by the color in the corresponding panel on the top right. Pearson correlations also indicated in the top right panels. In the panels on the diagonal, the distribution of scores for a given condition and replicate is shown. The replicate number is indicated above and to the right of the plots.

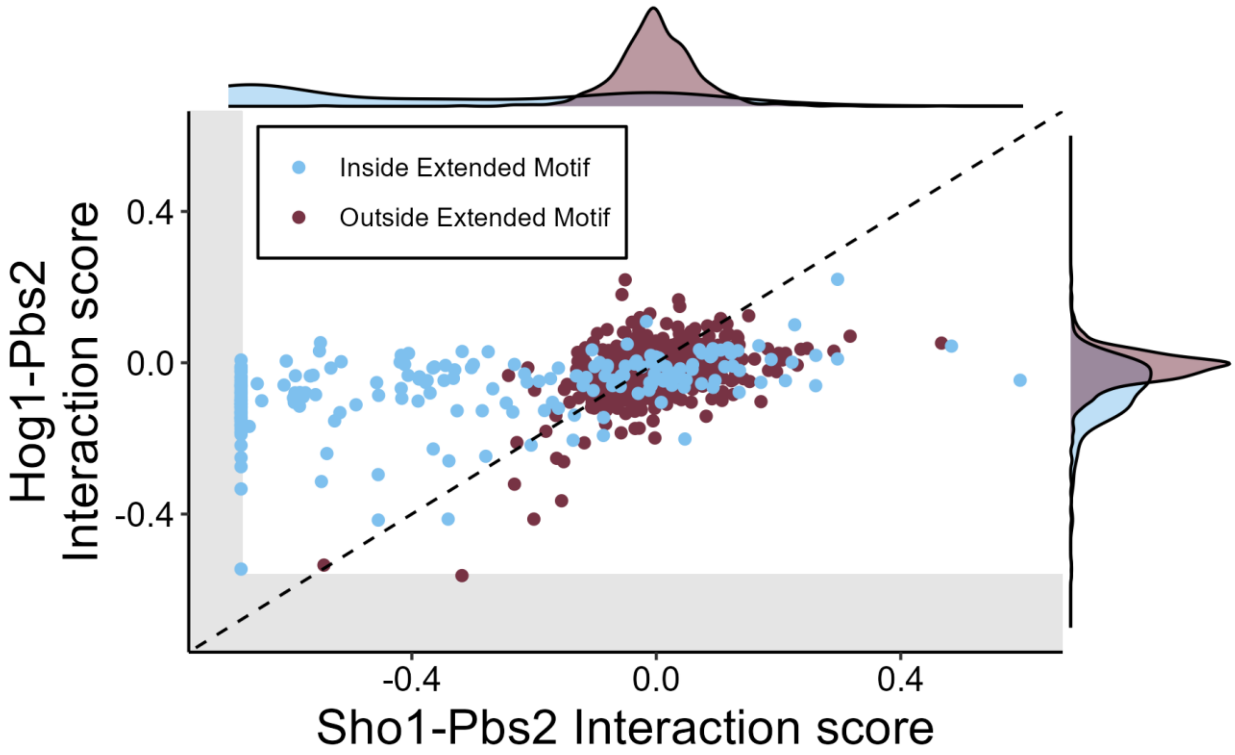


Figure S3.
Scatterplot of interaction scores for missense and silent mutants in the DMS library of the region surrounding the motif (positions 71-126), for the interaction of Pbs2 with either Sho1 or Hog1. Points are colored according to whether the mutants are situated inside or outside the extended motif (positions 85-99). The dotted line represents the diagonal. The gray rectangles on the left and bottom of the panel represent the 97.5 percentile of scores for the nonsense variants of Pbs2, which do not express the DHFR F[3] fragment. As such, they represent the limit of detectable signal in the assay. The points which have scores comparable to the nonsense mutations, and therefore no detectable interaction, were placed at the limit of the gray rectangle to indicate that no interaction was detected. In the margins, density plots show the distribution of scores for mutants situated either inside or outside the extended motif (85-99). Each point represents the median of all codons in each replicate coding for the same mutation, with 3 to 18 replicates for each mutant.

**a** Sho1

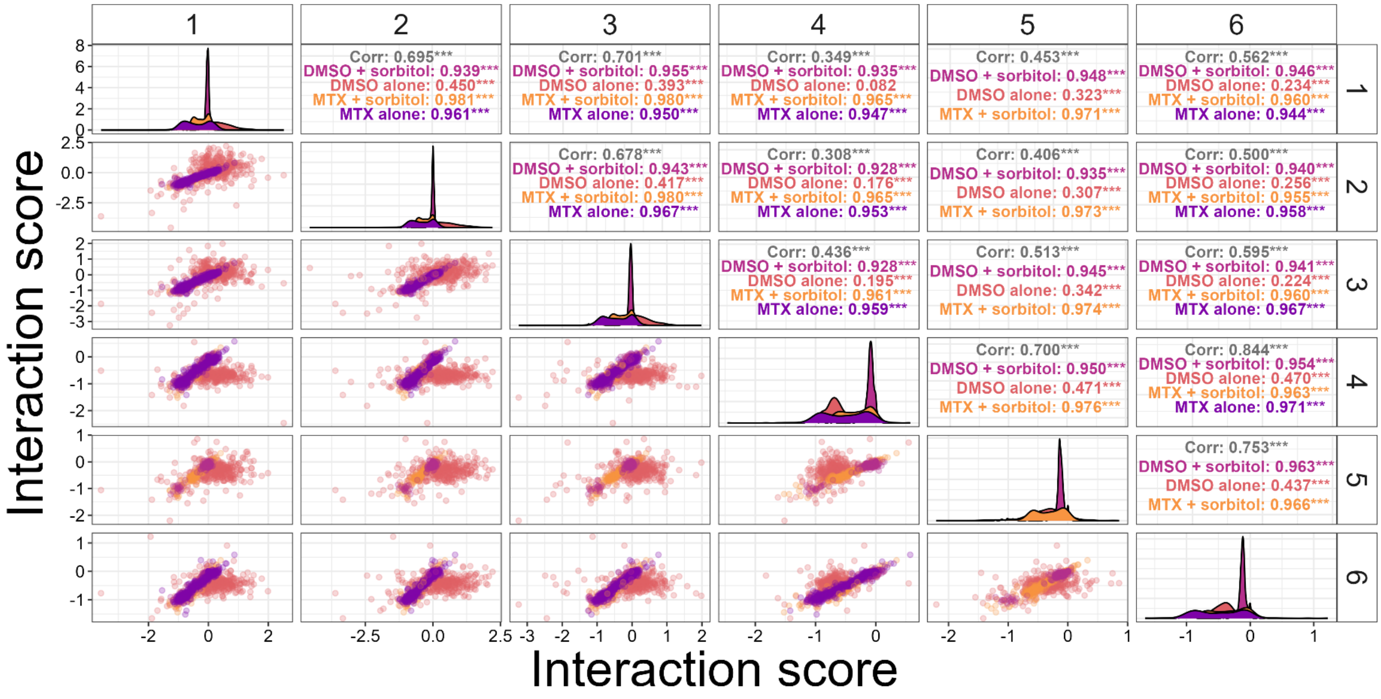

1
2
3
4
5
6
Corr: 0.695***
DMSO + sorbitol: 0.939***
DMSO alone: 0.450***
MTX + sorbitol: 0.981***
MTX alone: 0.961***
Corr: 0.701***
DMSO + sorbitol: 0.955***
DMSO alone: 0.393***
MTX + sorbitol: 0.980***
MTX alone: 0.950***
Corr: 0.349***
DMSO + sorbitol: 0.935***
DMSO alone: 0.082
MTX + sorbitol: 0.965***
MTX alone: 0.947***
Corr: 0.453***
DMSO + sorbitol: 0.948***
DMSO alone: 0.323***
MTX + sorbitol: 0.971***
Corr: 0.562***
DMSO + sorbitol: 0.946***
DMSO alone: 0.234***
MTX + sorbitol: 0.960***
MTX alone: 0.944***
Corr: 0.678***
DMSO + sorbitol: 0.943***
DMSO alone: 0.417***
MTX + sorbitol: 0.980***
MTX alone: 0.967***
Corr: 0.308***
DMSO + sorbitol: 0.928***
DMSO alone: 0.176***
MTX + sorbitol: 0.965***
MTX alone: 0.953***
Corr: 0.406***
DMSO + sorbitol: 0.935***
DMSO alone: 0.307***
MTX + sorbitol: 0.973***
Corr: 0.500***
DMSO + sorbitol: 0.940***
DMSO alone: 0.256***
MTX + sorbitol: 0.955***
MTX alone: 0.958***
Corr: 0.436***
DMSO + sorbitol: 0.928***
DMSO alone: 0.195***
MTX + sorbitol: 0.961***
MTX alone: 0.959***
Corr: 0.513***
DMSO + sorbitol: 0.945***
DMSO alone: 0.342***
MTX + sorbitol: 0.974***
Corr: 0.595***
DMSO + sorbitol: 0.941***
DMSO alone: 0.224***
MTX + sorbitol: 0.960***
MTX alone: 0.967***
Corr: 0.700***
DMSO + sorbitol: 0.950***
DMSO alone: 0.471***
MTX + sorbitol: 0.976***
Corr: 0.844***
DMSO + sorbitol: 0.954***
DMSO alone: 0.470***
MTX + sorbitol: 0.963***
MTX alone: 0.971***
Corr: 0.753***
DMSO + sorbitol: 0.963***
DMSO alone: 0.437***
MTX + sorbitol: 0.966***
Interaction score
Interaction score


**b** Hog1

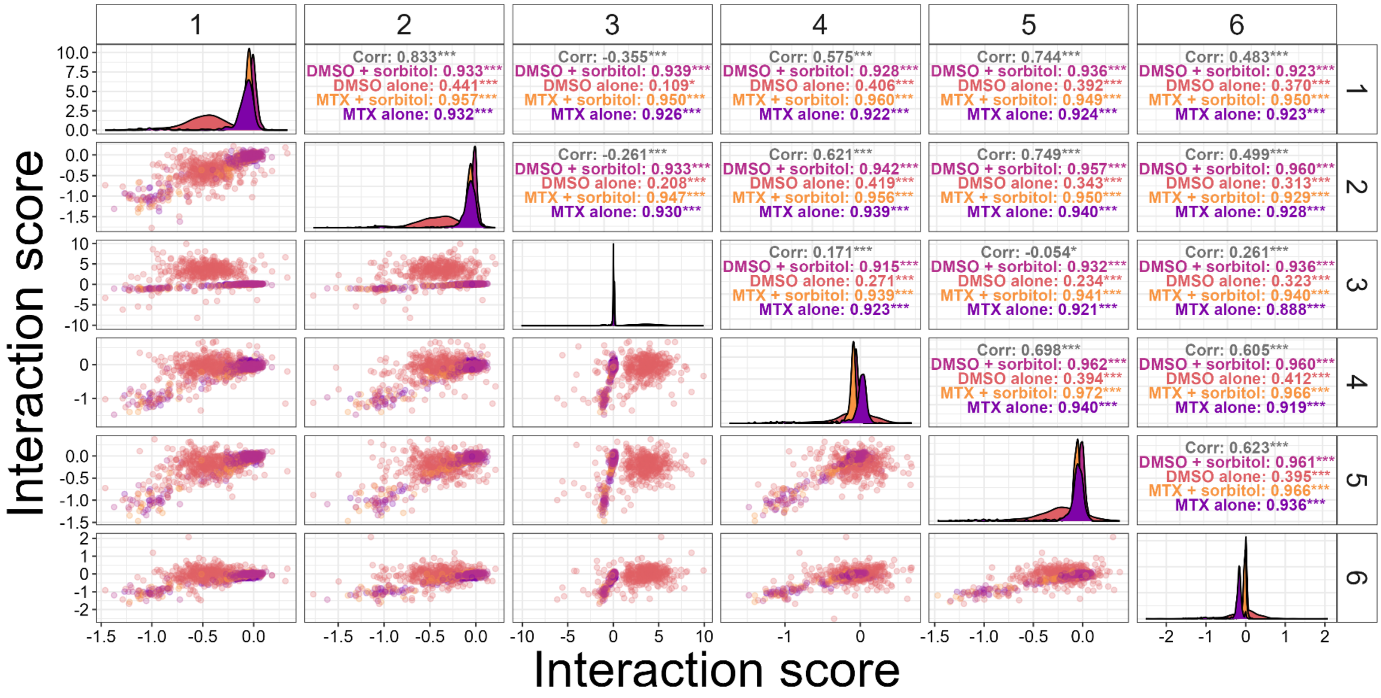

1
2
3
4
5
6
Corr: 0.833***
DMSO + sorbitol: 0.933***
DMSO alone: 0.441***
MTX + sorbitol: 0.957***
MTX alone: 0.932***
Corr: -0.355***
DMSO + sorbitol: 0.939***
DMSO alone: 0.109*
MTX + sorbitol: 0.950***
MTX alone: 0.926***
Corr: 0.575***
DMSO + sorbitol: 0.928***
DMSO alone: 0.406***
MTX + sorbitol: 0.960***
MTX alone: 0.922***
Corr: 0.744***
DMSO + sorbitol: 0.936***
DMSO alone: 0.392***
MTX + sorbitol: 0.949***
MTX alone: 0.924***
Corr: 0.483***
DMSO + sorbitol: 0.923***
DMSO alone: 0.370***
MTX + sorbitol: 0.950***
MTX alone: 0.923***
Corr: -0.261***
DMSO + sorbitol: 0.933***
DMSO alone: 0.208***
MTX + sorbitol: 0.947***
MTX alone: 0.930***
Corr: 0.621***
DMSO + sorbitol: 0.942***
DMSO alone: 0.419***
MTX + sorbitol: 0.956***
MTX alone: 0.939***
Corr: 0.749***
DMSO + sorbitol: 0.957***
DMSO alone: 0.343***
MTX + sorbitol: 0.950***
MTX alone: 0.940***
Corr: 0.499***
DMSO + sorbitol: 0.960***
DMSO alone: 0.313***
MTX + sorbitol: 0.929***
MTX alone: 0.928***
Corr: 0.171***
DMSO + sorbitol: 0.915***
DMSO alone: 0.271***
MTX + sorbitol: 0.939***
MTX alone: 0.923***
Corr: -0.054*
DMSO + sorbitol: 0.932***
DMSO alone: 0.234***
MTX + sorbitol: 0.941***
MTX alone: 0.921***
Corr: 0.261***
DMSO + sorbitol: 0.936***
DMSO alone: 0.323***
MTX + sorbitol: 0.940***
MTX alone: 0.888***
Corr: 0.698***
DMSO + sorbitol: 0.962***
DMSO alone: 0.394***
MTX + sorbitol: 0.972***
MTX alone: 0.940***
Corr: 0.605***
DMSO + sorbitol: 0.960***
DMSO alone: 0.412***
MTX + sorbitol: 0.966***
MTX alone: 0.919***
Corr: 0.623***
DMSO + sorbitol: 0.961***
DMSO alone: 0.395***
MTX + sorbitol: 0.966***
MTX alone: 0.936***
Interaction score
Interaction score

Figure S4.

<u>Correlation between replicates in the DHFR-PCA experiments on DMS libraries of the extended Pbs2 binding motif</u> a) Scatterplot between the 6 replicates of the Sho1-Pbs2 DHFR-PCA interaction scores for mutants in the DMS library of the extended Pbs2 motif (positions 85-100). Each point represents one Pbs2 codon variant in one condition (with or without sorbitol, with MTX or with only DMSO), as indicated by the color in the corresponding panel on the top right. Pearson correlations also indicated in the top right panels. In the panels on the diagonal, the distribution of scores for a given condition and replicate is shown. The replicate number is indicated above and to the right of the plots. b) Scatterplot between the 6 replicates of the Hog1-Pbs2 DHFR-PCA interaction scores for mutants in the DMS library of the extended Pbs2 motif (positions 85-100). Each point represents one Pbs2 codon variant in one condition (with or without sorbitol, with MTX or with only DMSO), as indicated by the color in the corresponding panel on the top right. Pearson correlations also indicated in the top right panels. In the panels on the diagonal, the distribution of scores for a given condition and replicate is shown. The replicate number is indicated above and to the right of the plots.

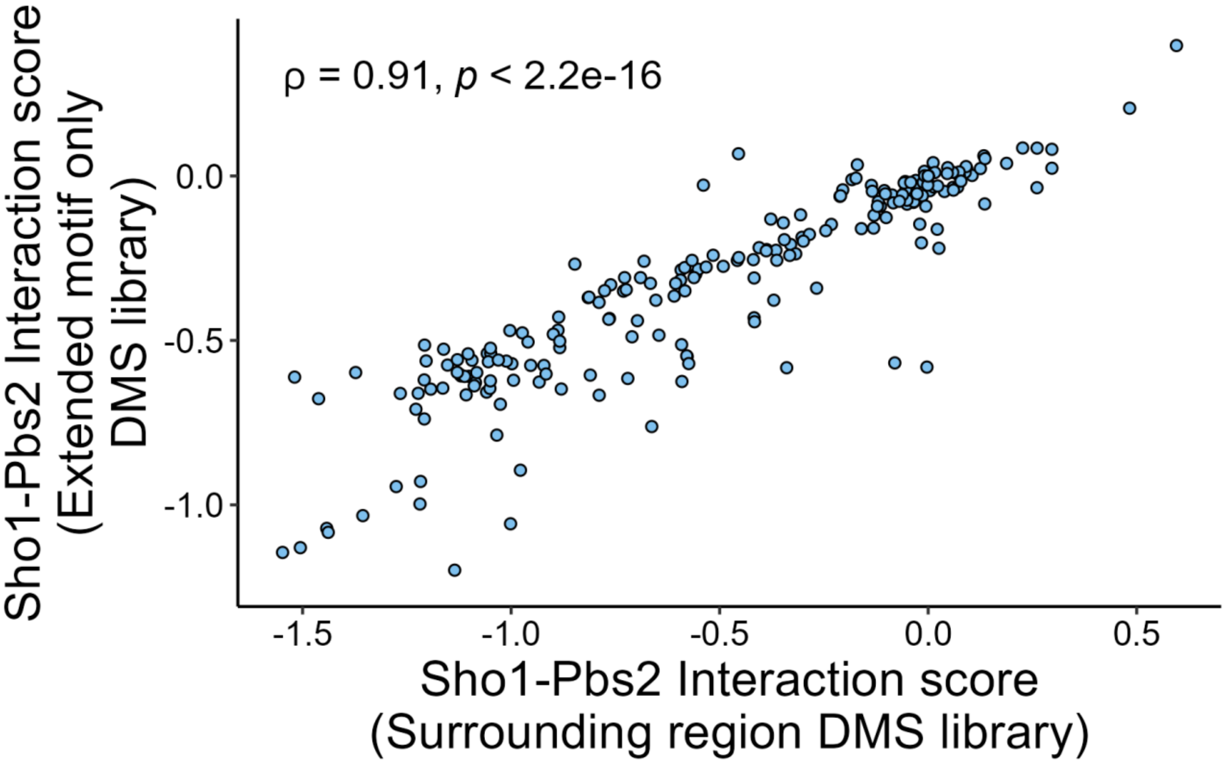


Figure S5.

Scatterplot of interaction scores of Pbs2 mutants in the initial DHFR-PCA screen on the DMS library of the surrounding region of the motif (positions 71-126), and the same mutants' interaction scores in the subsequent DHFR-PCA screen on the DMS library of only the extended motif (positions 85-100). Scores shown were measured in the presence of methotrexate and 1 M of sorbitol. Each point represents the median of all codons in each replicate coding for the same mutation, with 3 to 18 replicates for each mutant. Spearman's rho 0.91, $p < 2.2X10^{-16}$.

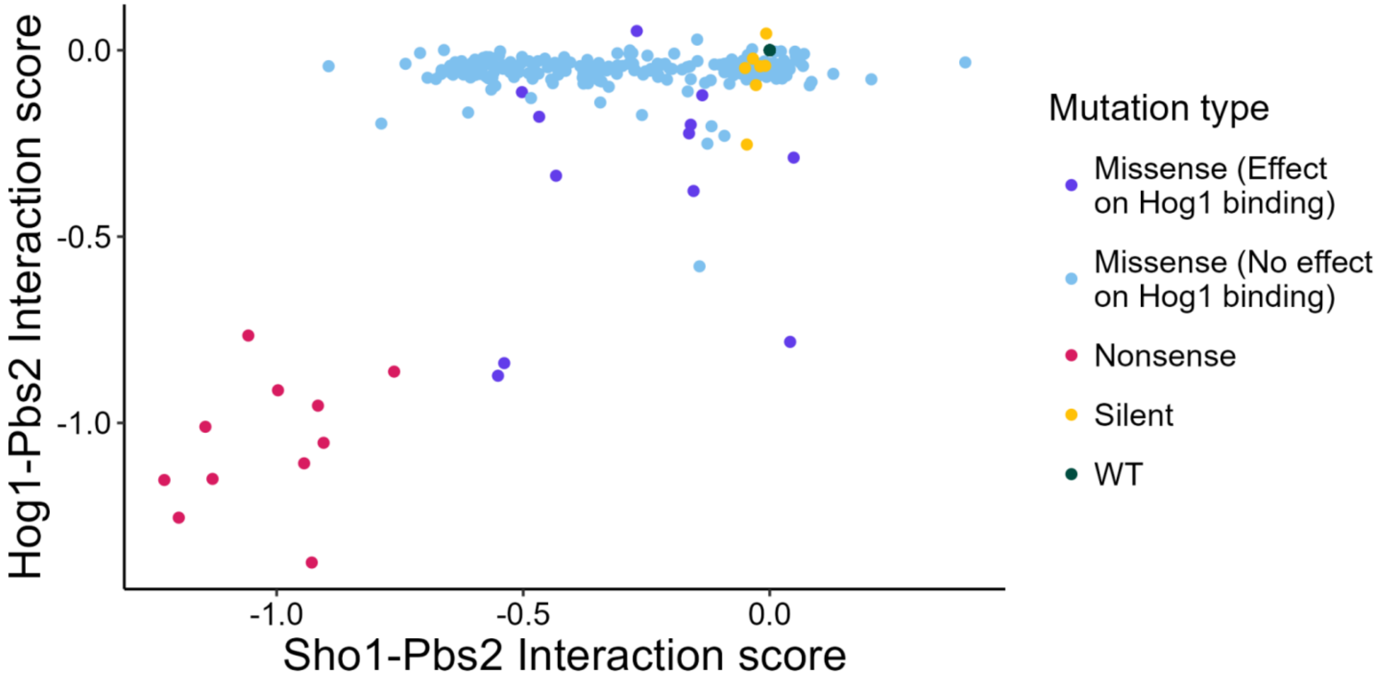


Figure S6.
Scatterplot of interaction scores of the Pbs2 extended motif DMS library (positions 85-100) mutants with Sho1 and Hog1, in the presence of 1 M of sorbitol. Purple missense points represent Pbs2 mutants which have an interaction score with Hog1 which is significantly different to the combination of silent mutants and wild-type Pbs2 (Two-sided Mann-Whitney U test with false discovery rate corrected p-value < 0.05), while blue missense points represent mutants for which the Hog1 interaction score is not significantly different to the combination of silent mutants and wild-type Pbs2. The combination of silent mutations and wild type represents all mutants expressing the wild-type protein sequence. Since the Hog1 interaction is used to measure abundance and stability of the Pbs2 mutants, the significantly different mutants are considered to have an effect on the abundance or the stability of Pbs2 and were not kept for the remainder of the analysis. The only exceptions are the nonsense mutants, which were kept because they provide a useful reference. Each point represents the median of all codons in each replicate coding for the same mutation, with 3 to 18 replicates for each mutant.

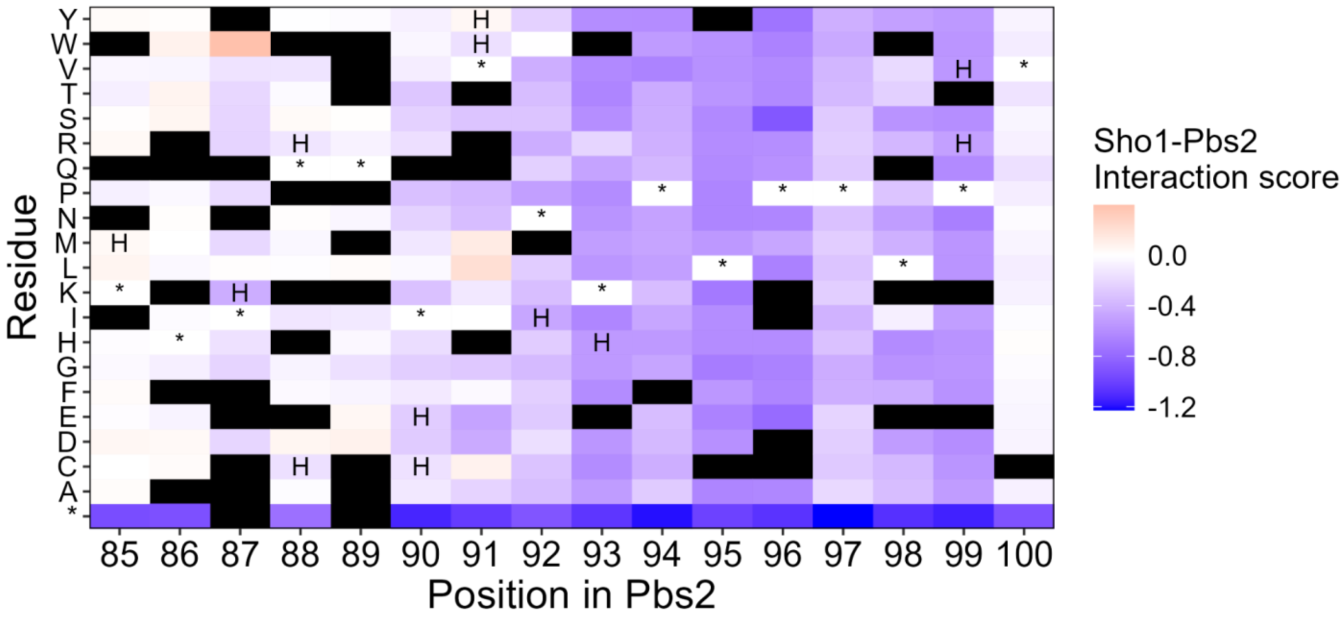


Figure S7.

Sho1-Pbs2 interaction scores for Pbs2 mutants in the DHFR-PCA on the DMS library of the extended Pbs2 motif (positions 85-100). Scores are indicated for each mutant according to their position in Pbs2 and the residue that it was mutated to. The wild-type residue for each position is indicated by an asterisk (*). Mutants which significantly affect the Hog1-Pbs2 interaction are marked with the letter H (Two-sided Mann-Whitney U test with false discovery rate corrected p-value < 0.05). Mutants which were removed from the dataset for having too few reads in the initial timepoint are marked in black. Each measurement represents the median of all codons in each replicate coding for the same residue, with 3 to 18 replicates for each mutant.

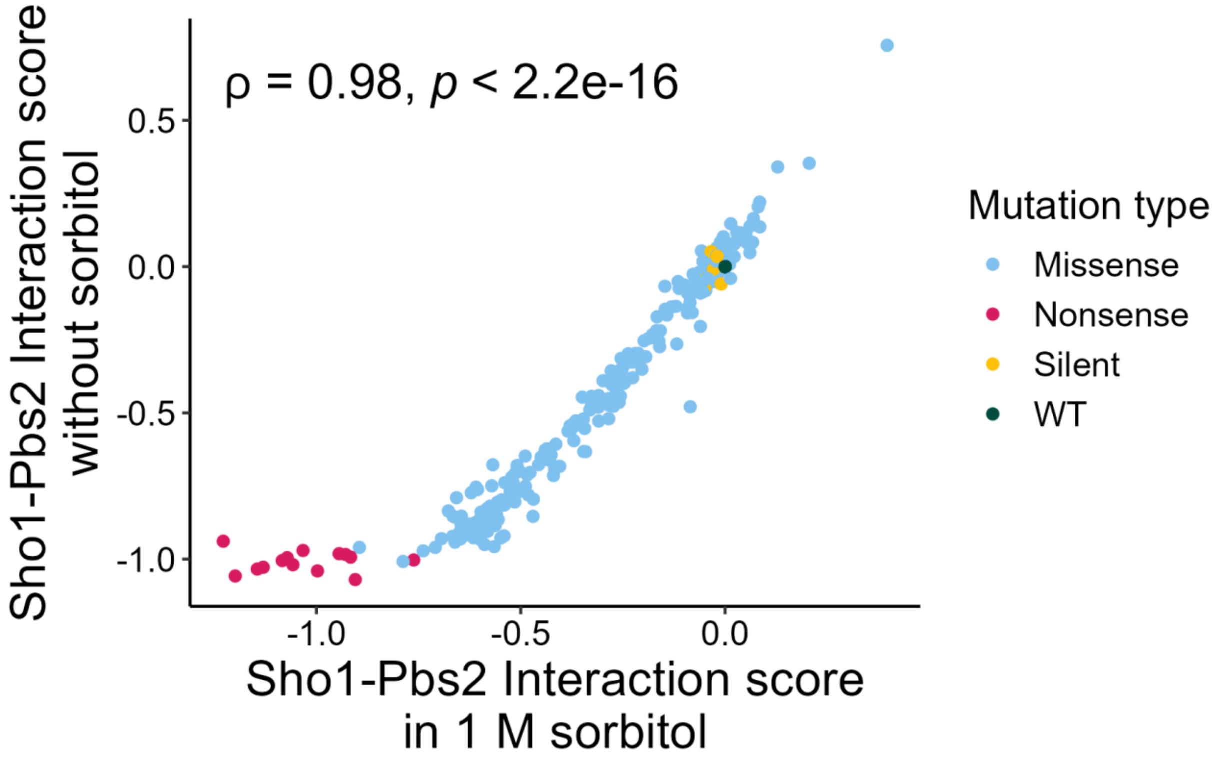


Figure S8.
Scatterplot of the Sho1 interaction scores for all mutants in the Pbs2 extended motif DMS library (positions 85-100), in the presence and absence of 1 M of sorbitol. Measurements in DHFR-PCA media with methotrexate. The correlation has a spearman's rho of 0.98, and a p-value < $2.2X10^{-16}$. The measurements for the interaction in 1 M sorbitol have a greater range due to the greater activation of the HOG pathway. Each point represents the median of all codons in each replicate coding for the same mutation, with 3 to 18 replicates for each mutant.

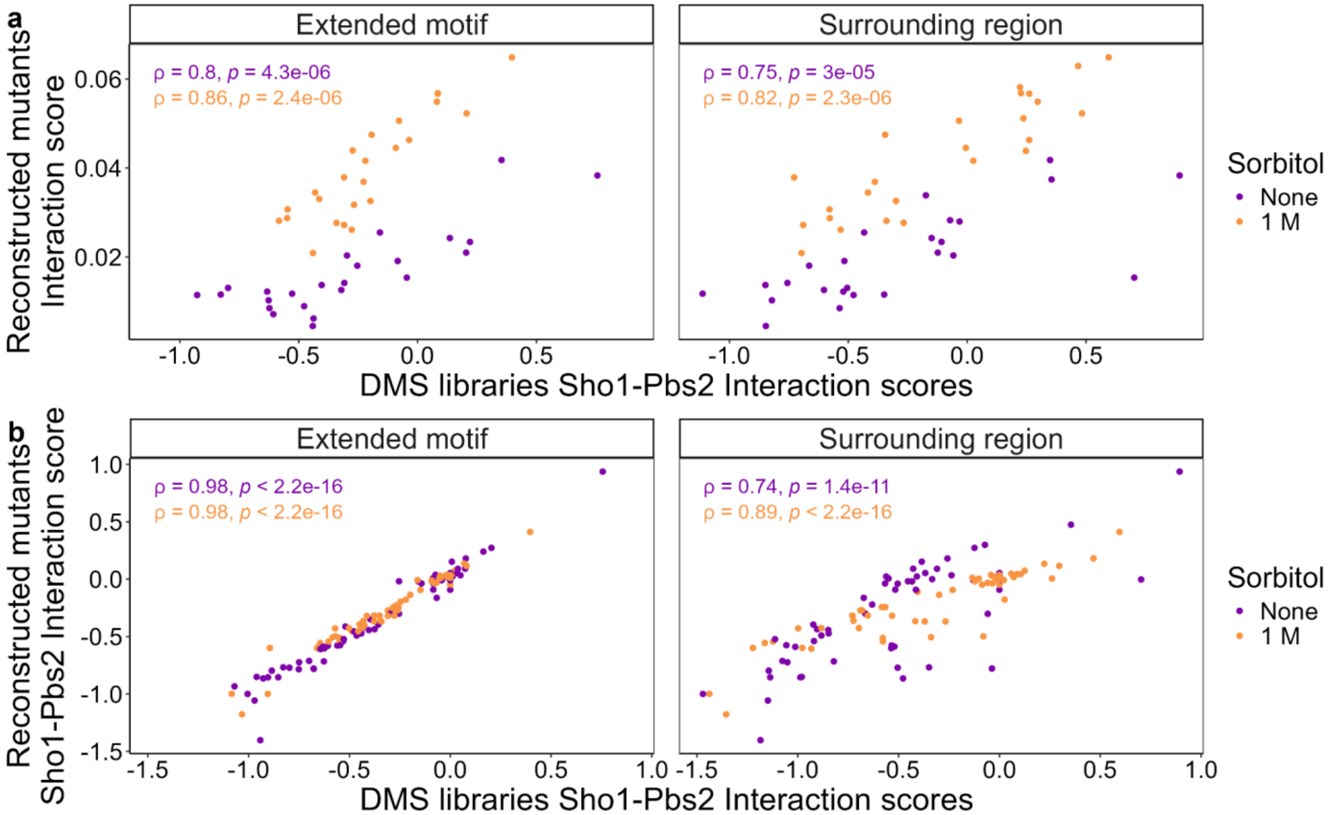


Figure S9.

<u>Validation of pooled DHFR-PCA effects by individually reconstructed mutants</u>

a) Scatterplot of Sho1-Pbs2 interaction scores as measured in pooled competition DHFR-PCA of the DMS libraries (x-axis) and growth rate of the same mutations individually reconstructed, in DHFR-PCA growth curves (y-axis). Results from the DMS library of the extended motif (positions 85-100) (left) and the DMS library of the surrounding region (positions 71-126) (right), in the presence (orange) or absence (purple) of 1 M of sorbitol. The spearman correlations between the two methods of measuring the interaction strength are indicated on the plots. The values for the DMS library interaction scores represents the median of all codons in each replicate coding for the same mutation, with 3 to 18 replicates for each mutant. The values for the growth rate represent the median of 3 replicates. b) Scatterplot of Sho1-Pbs2 interaction scores as measured in pooled competition DHFR-PCA of the DMS libraries (x-axis) and measured in pooled competition DHFR-PCA of individually reconstructed mutants (y-axis). Results from the DMS library of the extended motif (positions 85-100) (left) and the DMS library of the surrounding region (positions 71-126) (right), in the presence (orange) or absence (purple) of 1 M of sorbitol. The spearman correlations between the two methods of measuring the interaction strength are indicated on the plots. The values for the DMS library interaction scores represents the median of all codons in each replicate coding for the same mutation, with 3 to 18 replicates for each mutant. The values for the reconstructed mutants represent the median of 3 to 6 replicates.

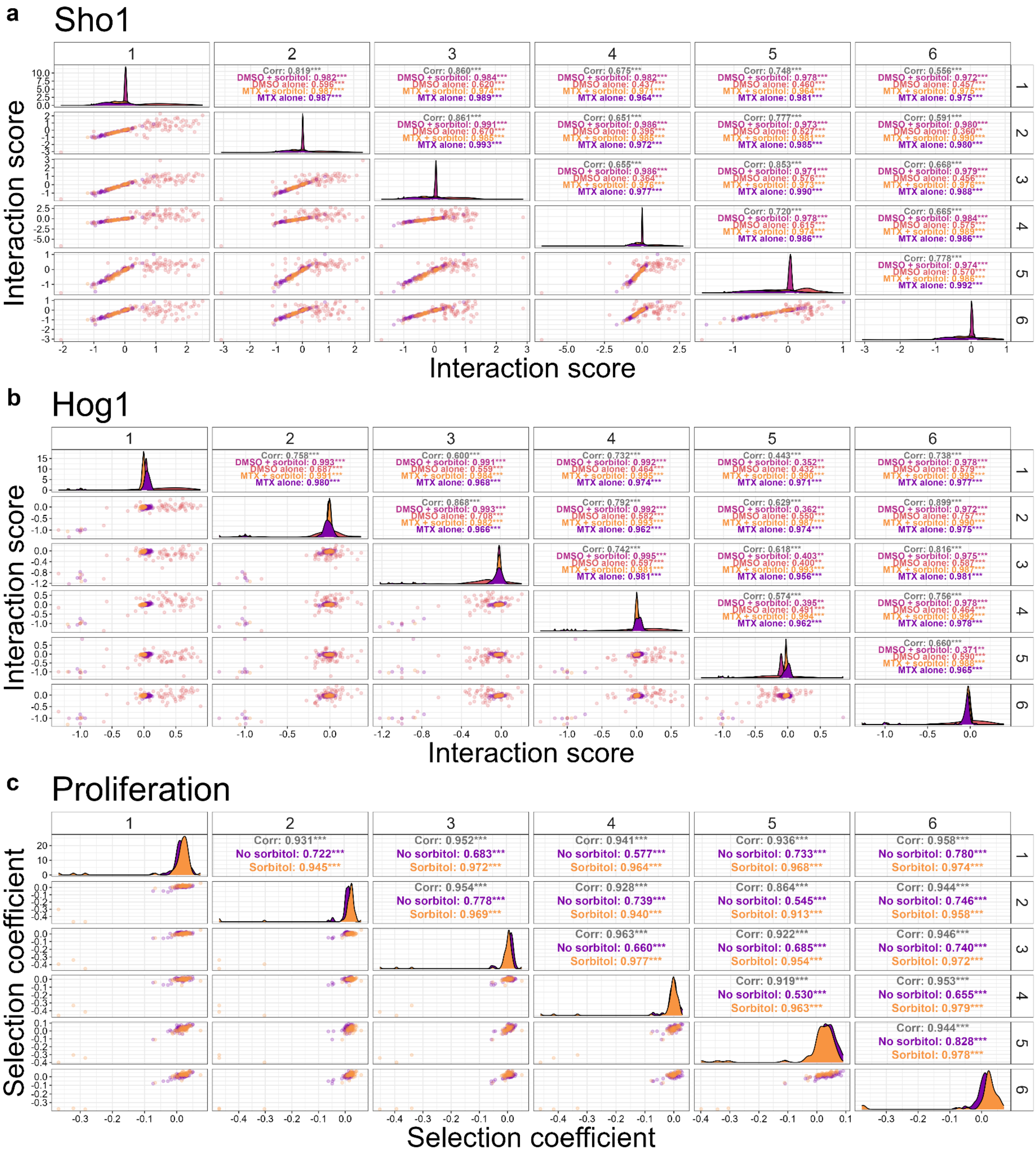
a
Sho1
Interaction score
Interaction score
b
Hog1
Interaction score
Interaction score
c
Proliferation
Selection coefficient
Selection coefficient
Corr: 0.931***
No sorbitol: 0.722***
Sorbitol: 0.945***
Corr: 0.952***
No sorbitol: 0.683***
Sorbitol: 0.972***
Corr: 0.941***
No sorbitol: 0.577***
Sorbitol: 0.964***
Corr: 0.936***
No sorbitol: 0.733***
Sorbitol: 0.968***
Corr: 0.958***
No sorbitol: 0.780***
Sorbitol: 0.974***
Corr: 0.954***
No sorbitol: 0.778***
Sorbitol: 0.969***
Corr: 0.928***
No sorbitol: 0.739***
Sorbitol: 0.940***
Corr: 0.864***
No sorbitol: 0.545***
Sorbitol: 0.913***
Corr: 0.944***
No sorbitol: 0.746***
Sorbitol: 0.958***
Corr: 0.963***
No sorbitol: 0.660***
Sorbitol: 0.977***
Corr: 0.922***
No sorbitol: 0.685***
Sorbitol: 0.954***
Corr: 0.946***
No sorbitol: 0.740***
Sorbitol: 0.972***
Corr: 0.919***
No sorbitol: 0.530***
Sorbitol: 0.963***
Corr: 0.953***
No sorbitol: 0.655***
Sorbitol: 0.979***
Corr: 0.944***
No sorbitol: 0.828***
Sorbitol: 0.978***

Figure S10.

<u>Correlation between replicates in the DHFR-PCA experiments on validation libraries of the extended Pbs2 binding motif</u>

a) Scatterplot between the 6 replicates of the Sho1-Pbs2 DHFR-PCA interaction scores for mutants in the Pbs2 validation library. Each point represents one Pbs2 codon variant in one condition (with or without sorbitol, with MTX or with only DMSO), as indicated by the color in the corresponding panel on the top right. Pearson correlations also indicated in the top right panels. In the panels on the diagonal, the distribution of scores for a given condition and replicate is shown. The replicate number is indicated above and to the right of the plots. b) Scatterplot between the 6 replicates of the Hog1-Pbs2 DHFR-PCA interaction scores for mutants in the Pbs2 validation library. Each point represents one Pbs2 codon variant in one condition (with or without sorbitol, with MTX or with only DMSO), as indicated by the color in the corresponding panel on the top right. Pearson correlations also indicated in the top right panels. In the panels on the diagonal, the distribution of scores for a given condition and replicate is shown. The replicate number is indicated above and to the right of the plots. c) Scatterplot between the 6 replicates of selection coefficients for mutants in the Pbs2 validation library measuring cell proliferation. Each point represents one Pbs2 codon variant in one condition (with or without sorbitol), as indicated by the color in the corresponding panel on the top right. Pearson correlations also indicated in the top right panels. In the panels on the diagonal, the distribution of scores for a given condition and replicate is shown. The replicate number is indicated above and to the right of the plots.

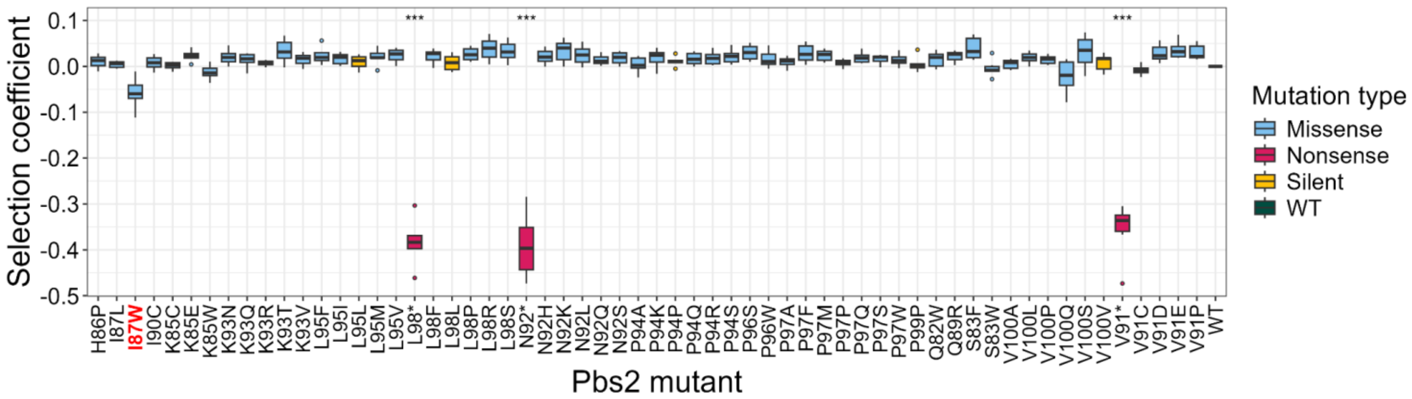


Figure S11.

Impact on growth rate measured as selection coefficients for individually reconstructed Pbs2 mutants in SC synthetic media with 1 M sorbitol. The strains contain no DHFR fragments on any proteins. Boxplots show the distribution of 6 replicates per mutant. The selection coefficient is normalized by the growth of the wild-type sequence, making the wild-type selection coefficient zero. Only the 3 nonsense mutants grew significantly less than wild-type Pbs2 and the silent mutants (Welch's t-test with false discovery rate correction p-value < 0.05), although I87W (highlighted in red on axis), which is the mutant with the highest Sho1-Pbs2 interaction score shows a marked decrease in selection coefficient (p-value = 0.066).

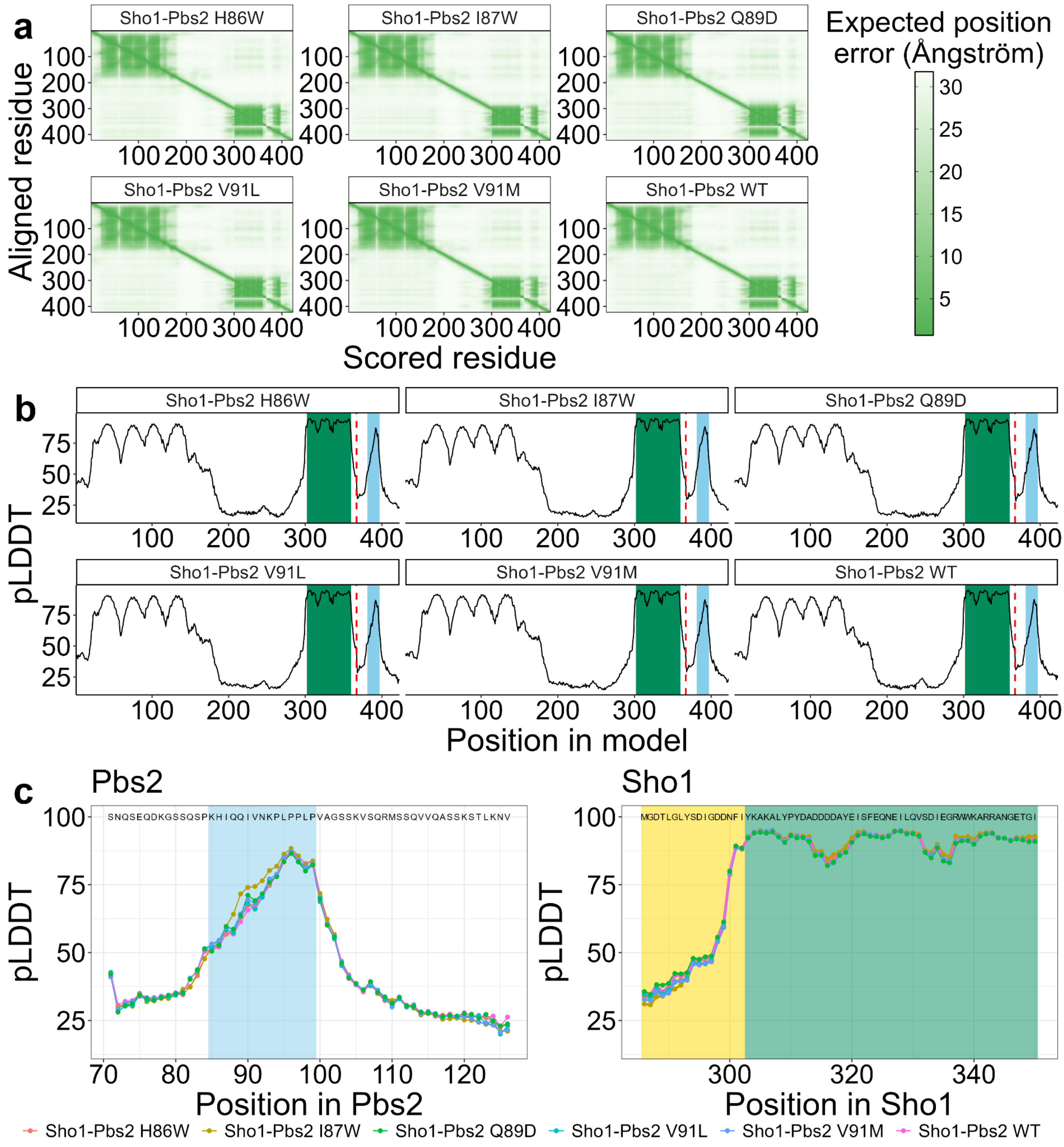
a
Sho1-Pbs2 H86W
Sho1-Pbs2 I87W
Sho1-Pbs2 Q89D
Sho1-Pbs2 V91L
Sho1-Pbs2 V91M
Sho1-Pbs2 WT
Aligned residue
Scored residue
Expected position error (Ångström)
b
pLDDT
Position in model
c
Pbs2
Sho1
SNQSEQDKGSSQSPKHIQQIVNKPLPPLPVAGSSKVSQRMSSQVVQASSKSTLKNV
MGDTLGLYSDIGDDNFIYKAKALYPYDADDDDAYEISFEQNEILQVSDIEGRWWKARRANGETGI
Position in Pbs2
Position in Sho1

Figure S12.

Confidence metrics for the seven structures predicted using AlphaFold2-Multimer. Sho1-Pbs2 WT indicates the structure in figure 3a and 3b, Sho1-Pbs2 I87W indicates the structure in figure 3c, and all others are the structures with the same Pbs2 mutants in Figure S13 a) Predicted aligned error (PAE) values for the 6 predicted structures, output from AlphaFold-Multimer. b) pLDDT value for each position in the 6 predicted structures, output from AlphaFold-Multimer. Positions corresponding to the SH3 domain (positions 303-360) of Sho1 are indicated by a green background, and positions corresponding to the extended motif (positions 85-99) of Pbs2 are indicated by a blue background. Since AlphaFold-Multimer combines the sequences of both proteins into a single output, a red dashed line was used to mark the end of the Sho1 portion of the model, and the beginning of the Pbs2 portion of the model. c) pLDDT value for each position, for selected regions of the 6 predicted structures. For Pbs2, the extended motif has a light blue background. For Sho1, the non-SH3 portion has a yellow background while the SH3 domain has a green background. The amino acid sequences of the selected regions of Sho1 and Pbs2 are shown at the top of the panels.

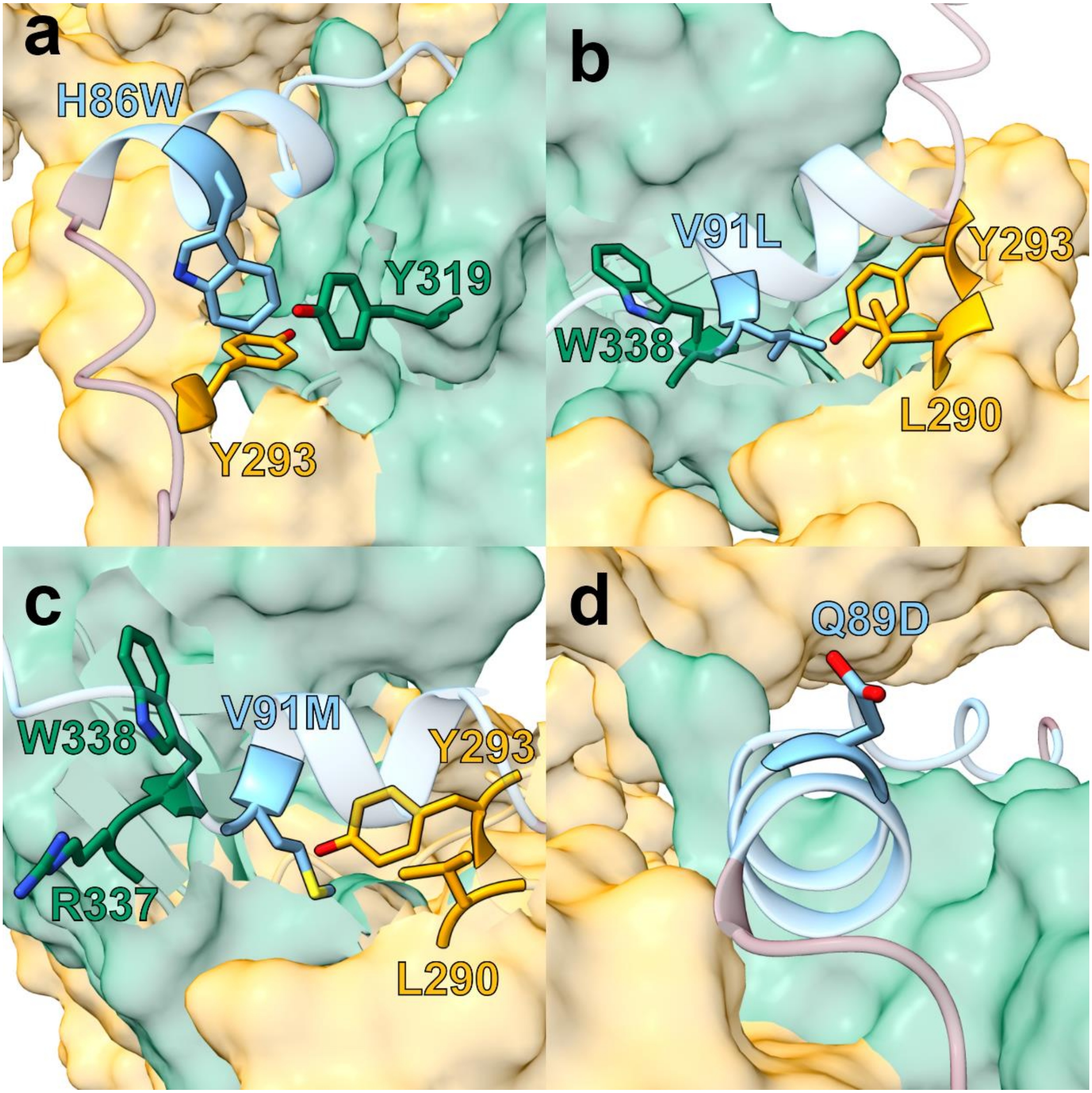


Figure S13.

Close-up details of AlphaFold-Multimer predicted complexes of Sho1 with Pbs2 mutants

Predicted structures of Sho1 in complex Pbs2 mutant a) H86W, b) V91L, c) V91M and d) Q89D. The predicted structures are colored as in panel a of figure 3, with the SH3 domain of Sho1 (position 303-360) in green, the non-SH3 portions of Sho1 in yellow, and the extended motif of Pbs2 (positions 85-99) in light blue, and the rest of Pbs2 in dark red. The mutated side chain in Pbs2 is shown, and the side chains of the residues that are contacted by the mutated residue are also shown. Q89D is not predicted to form any contacts.

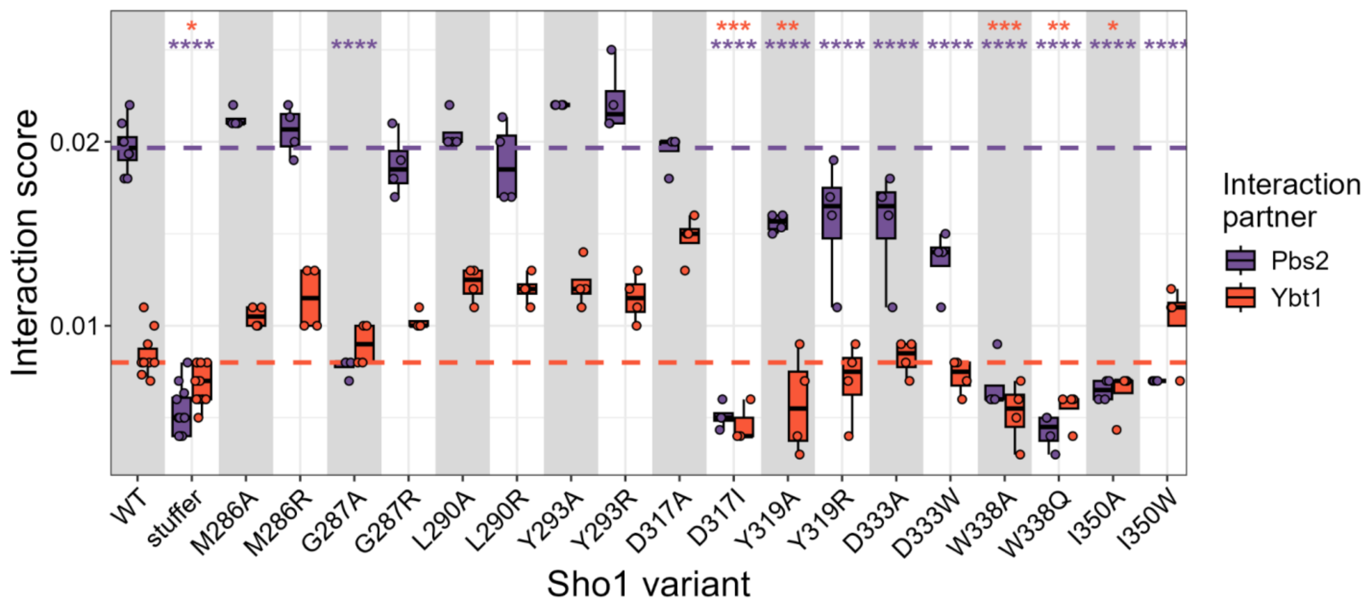


Figure S14.
Interaction scores of wild-type Sho1 and mutants with either Pbs2 or Ybt1, derived from growth rates from individual DHFR-PCA assays. Scores for individual replicates shown as points. Asterisks show the result of a one-sided Student's t-test with false discovery rate correction, comparing each mutant against the wild-type Sho1 with the same binding partner, and colored according to the partner. The median wild-type Sho1 interaction score with Pbs2 (purple) or Ybt1 (orange) is shown with dashed lines. Stuffer indicates a Sho1 mutant where the SH3 domain was replaced by a neutral, flexible peptide (GGSSGGGG).
* = p-value < 0.05, ** = p-value < 0.01, *** = p-value < 0.001, **** = p-value < 0.0001

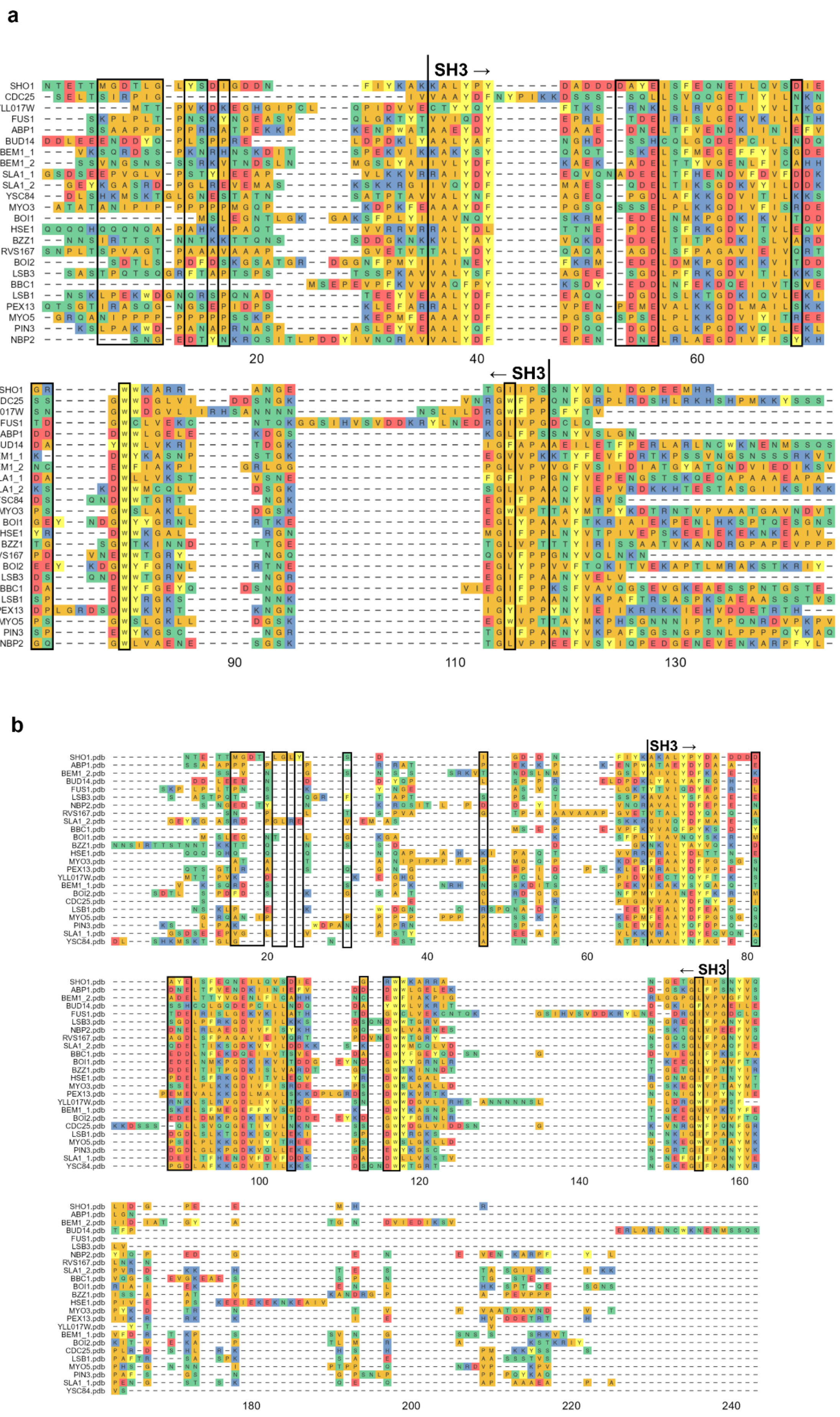
a
SH3 →
← SH3
b
SH3 →
← SH3

Figure S15.

Sequence and structural alignments of yeast SH3 domains and adjacent sequences a) Multiple sequence alignment of yeast SH3 domains with 25 residues on each side. Sho1 residues predicted to be within 5 Å of the non-canonical section of the Pbs2 extended motif (positions 85-92) are surrounded by a black rectangle. Beginning and end of the SH3 domain of Sho1 marked above the alignment. b) Multiple structural alignment of predicted yeast SH3 domains with 25 residues on each side, from MUSTANG. Residues which in the predicted Sho1-Pbs2 structure are within 5 Å of Pbs2 residues situated in the extended motif but not the canonical motif (positions 85-92) are surrounded by a black rectangle. Beginning and end of the SH3 domain of Sho1 marked above the alignment. Visualized using the R package ggmsa (Zhou *et al.* 2022).

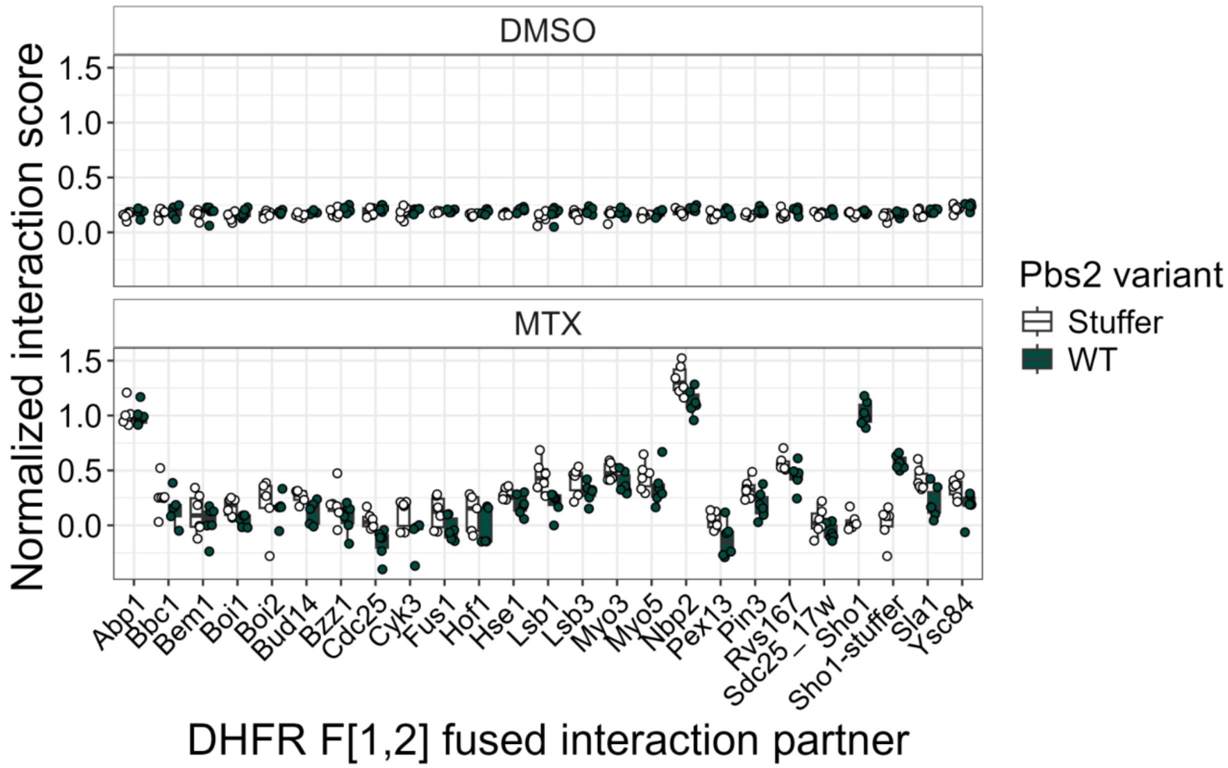


Figure S16.

Interaction score of wild-type Pbs2 (green) or Pbs2 where positions 76 to 126 have been replaced with a flexible stuffer sequence (Stuffer, white) with all yeast SH3-containing proteins. Controls grown on DMSO and not MTX, and therefore not measuring the interaction, are shown in the top panel. All replicates shown as points. Scores normalized with 1 being the median of Sho1 - wild-type Pbs2 interactions and 0 being the median of Sho1 - Pbs2-stuffer interactions. A control where the SH3 domain of Sho1 was replaced with the neutral, flexible stuffer sequence was also included, with the name Sho1-stuffer.

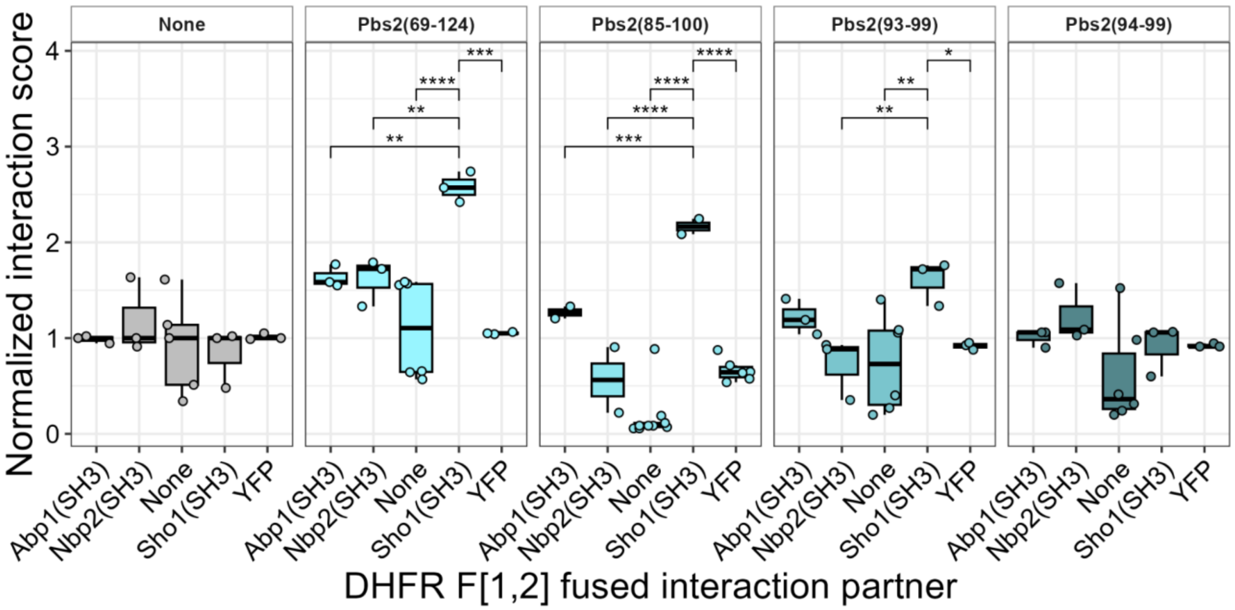


Figure S17.

Interaction scores of Pbs2 fragments, indicated in the strips above the plots, fused with DHFR F[3] with different SH3 domains, or a YFP control, fused with DHFR F[1,2]. “None” indicates that the DHFR fragment was expressed alone, with no SH3 domain or Pbs2 fragment fused to the fragment. SH3 domains were expressed under the *GAL1* promoter, and transcription was driven by an artificial transcription factor, inducible by β-estradiol, creating a system where expression of all SH3-DHFR F[1,2] constructs was at the same level (Aranda-Díaz *et al.* 2017). This removed the bias resulting from the unequal abundance of Abp1, Nbp2 and Sho1 such that the PCA signal does not reflect differences in abundance. For all growth assays, the expression of all SH3 domains was driven by 20 nM of β-estradiol, while the Pbs2 fragments were expressed from a plasmid with a constitutive promoter. The interaction scores are derived from growth rates, and normalized for each protein by the median interaction of the same protein against the DHFR F[3] attached to no Pbs2 fragment. The Pbs2(85-100) fragment has reduced availability for binding, as indicated by its relatively lower YFP-DHFR F[1,2] interaction score. This could be a result of increased degradation, or sequestration by Sho1. However, the relative interaction scores of Pbs2(85-100) with the other partners still clearly show a preference for Sho1 binding. Asterisks show the results of pairwise one-sided Student’s t-tests with false discovery rate corrected p-values, comparing the Sho1(SH3) interaction scores to the others.

* = p-value < 0.05, ** = p-value < 0.01, *** = p-value < 0.001, **** = p-value < 0.0001

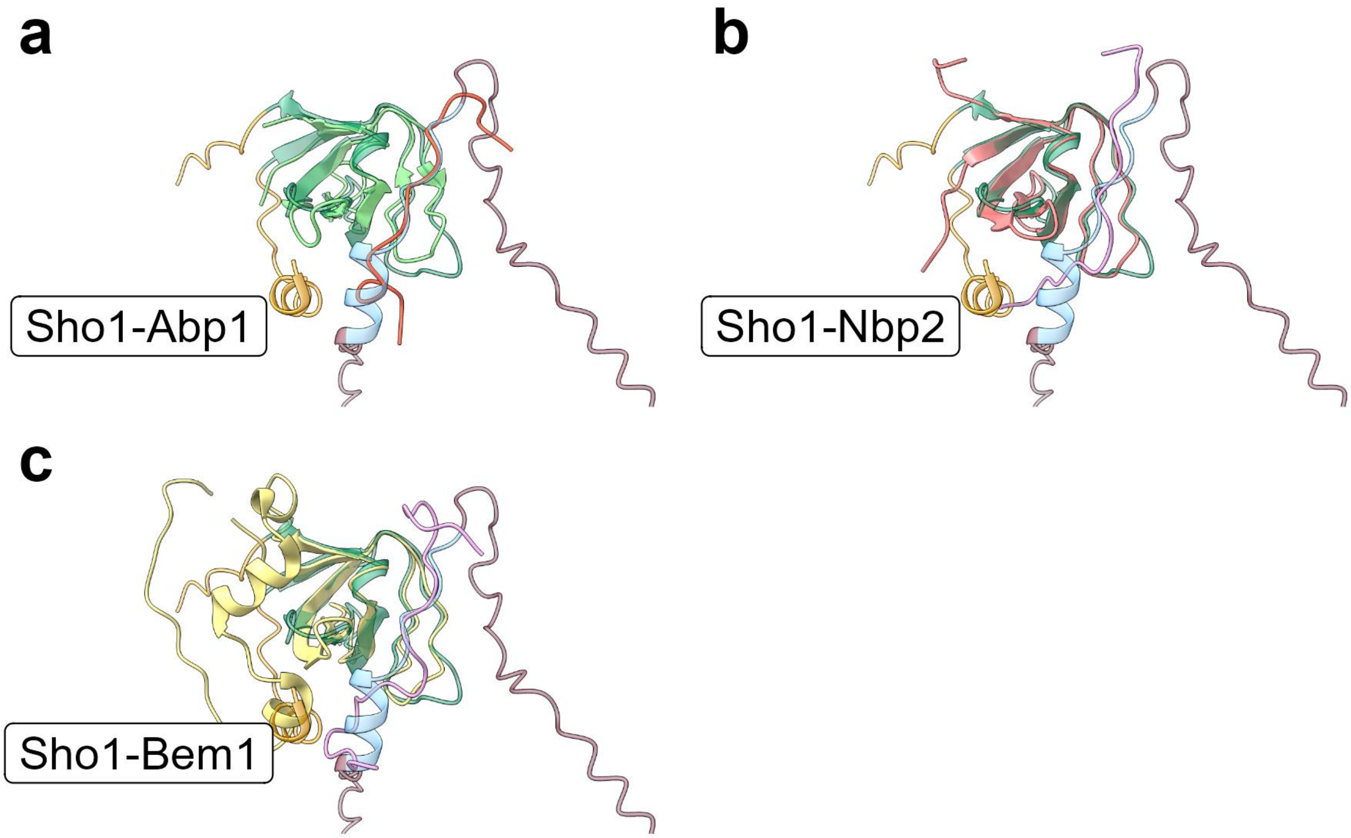


Figure S18.
Predicted structure of the Sho1-Pbs2 interaction superimposed with other SH3-motif pairs. The Sho1-Pbs2 structure is semi-transparent and colored as in figure 3a, with the SH3 domain of Sho1 (positions 303-360) in green, the non-SH3 portions of Sho1 in yellow, the Pbs2 extended motif (positions 85-99) in light blue and the rest of Pbs2 in dark red. Sho1-Pbs2 structure was superimposed with the a) Abp1-Ark structure (Abp1 - light green, Ark1 - pink), b) Nbp2-Ste20 structure (Nbp2 - orange-pink, Ste20 - light pink), and c) Bem1-2-Ste20 structure (Bem1 - yellow, Ste20 - light pink). Structures were superimposed using the Chimera X tool “Matchmaker” (Pettersen *et al.* 2021) to minimize root mean square deviance (RMSD) of backbone atom distances of the SH3 domains with Sho1.

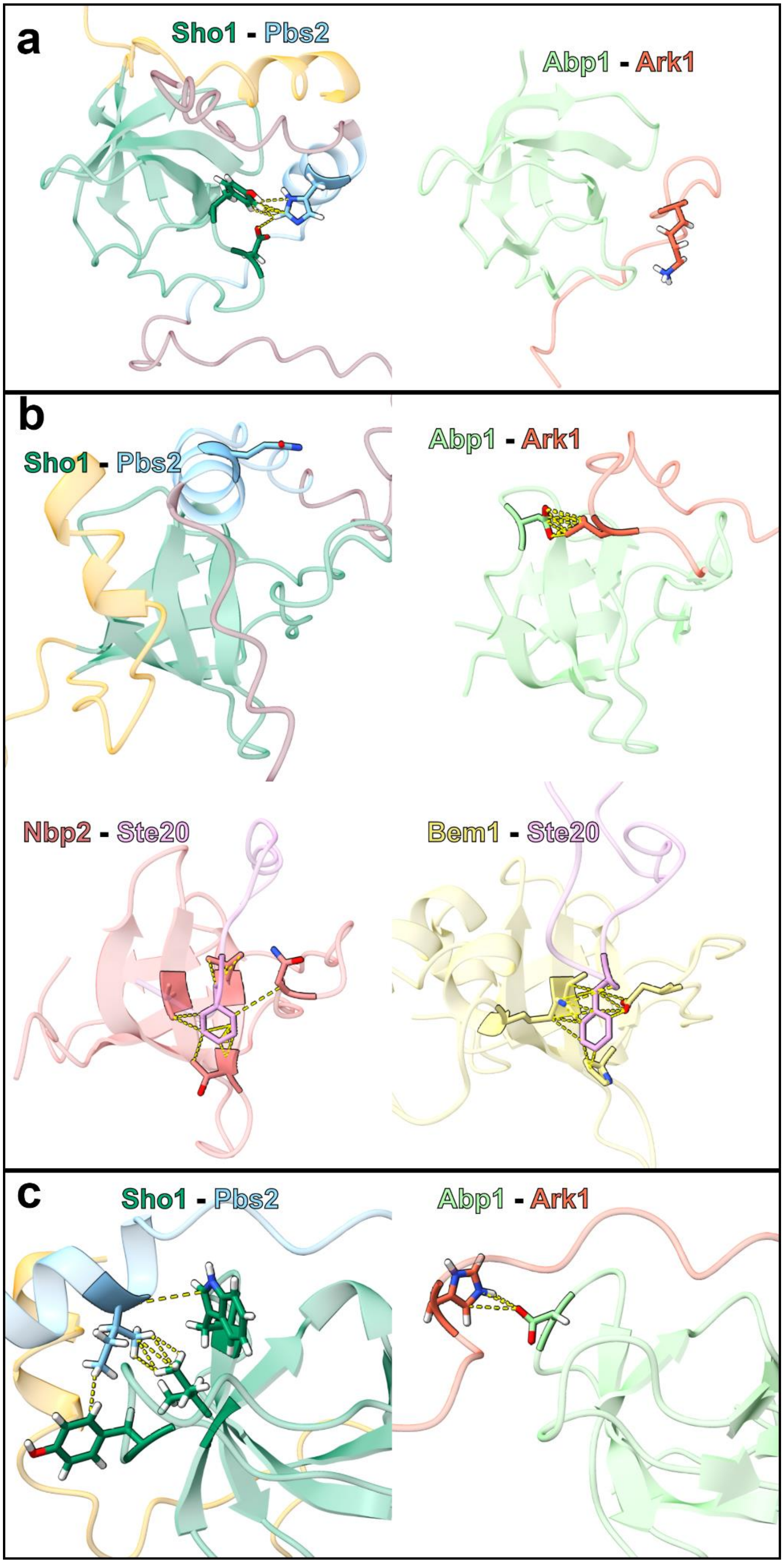
a
Sho1 - Pbs2
Abp1 - Ark1
b
Sho1 - Pbs2
Abp1 - Ark1
Nbp2 - Ste20
Bem1 - Ste20
c
Sho1 - Pbs2
Abp1 - Ark1

Figure S19.

<u>Differing contacts between SH3-extended motif pairs</u>

Previously measured NMR structures of Abp1-Ark1 (PDB accession 2RPN), Nbp2-Ste20 structure (PDB accession 2LCS) and Bem1-Ste20 (PDB accession 2RQW) were aligned to the predicted Sho1-Pbs2 structure using the ChimeraX tool “Matchmaker” (Pettersen *et al.* 2021), minimizing the RMSD between backbone atoms of the SH3 domains. The Sho1-Pbs2 predicted structure is colored as in figure 3a, with the SH3 domain of Sho1 (positions 303-360) in green, the non-SH3 portions of Sho1 in yellow, the Pbs2 extended motif (positions 85-99) in light blue and the rest of Pbs2 in dark red. Contacts between atoms are shown with dashed yellow lines. a) The side chains of positions -10 of Pbs2 and Ark1 and their contacts are shown, highlighting that the interaction made by Pbs2 is not present in Ark1. b) The side chains of position -7 of all extended motifs and their contacts are shown. While Ark1 and Ste20 form contacts with their respective partners, the glutamine in Pbs2 is not predicted to form any contact. Hydrogen atoms omitted to clarify interpretation of structures. c) The side chains of position -6 of Pbs2 and Ark1 and their contacts are shown. Ark1 forms a contact with an aspartic acid position that is not present in Sho1, and so the contacts formed by Pbs2 are different.
