## Supplementary File 3 - Supplementary Methods for "Residues Neighboring an SH3-Binding Motif Participate in the Interaction *In Vivo*"

Construction of DMS library

During the construction of the DMS library of the region surrounding the motif (codons 71 to 126 of *PBS2),* after the sequencing of the masterpool, three codons had unsatisfactory diversity. A new template was created for codons 103 and 119 using fusion PCR, creating a sequence of 967 base pairs around the mutated codon, using the original degenerate primers as well as new reverse degenerate primers to create an NNN codon at either codon 103 or 119. For codon 88, a new template was ordered as an oligonucleotide of 503 base pairs (Twist Bioscience), around codon 88, which is replaced by an NNN codon. These templates were transformed into Pbs2-stuffer-DHFR F[3] as described in the main text, and diversity at the desired codon was verified with Sanger sequencing. The three supplementary cultures were pooled into the masterpool, with an OD corresponding to 1/56th of the OD of the master pool.

Preparation of DNA sequencing libraries

The extracted DNA was amplified and barcoded using a Row-Column DNA barcoding strategy as previously described (Dubé *et al.* 2022). From the extracted genomic DNA, the *PBS2* region of interest was amplified using primers containing 3' overhangs which allow a second amplification to add barcodes. PCR products were diluted 1/2500, and used as a template to add row and column barcode primers onto the sequence, with a unique combination of 5' and 3' barcodes for each replicate in each condition. The row-column barcode primers were previously described (Dubé *et al.* 2022). PCR products were migrated on an agarose gel, and DNA concentrations were estimated based on band size using the Image Lab software (BioRad Laboratories). Samples were then pooled with different volumes to obtain the same quantity of DNA from each sample. The pools were purified using magnetic beads, measured using a NanoDrop (ThermoFisher), and diluted to 0.1 ng/µL. The diluted pools were then amplified with primers adding a second set of barcodes, called plate barcodes (Dubé *et al.* 2022). The sequencing of the first DHFR-PCA, on the larger region surrounding the Pbs2 motif (positions 70-126) was done using Illumina NovaSeq paired-end 250 base pair technology (CHUL sequencing platform, Quebec, Canada), while all other sequencing was done using Illumina MiSeq paired-end 300 bp technology (IBIS Genomic Analysis Platform, Quebec, Canada).

Visualization of DNA sequencing results

For the visualization of DMS DHFR-PCA results, ggplot2 (Wickham 2016) was used along with other R packages. The ggExtra (Attali and Baker 2023) command "ggMarginal" was used to add density plots to the exterior of Figure S3 (see File S1). The GGally (Schloerke *et al.* 2023) command "ggpairs" was used to make Figures S2, S4 and S10 (see File S1).

Analysis of contacts in SH3-motif structures

Contacts were analyzed for 4 structures. Sho1-Pbs2 was generated by AlphaFold-Multimer, as described in the main text. Abp1-Ark (PDB accession 2RPN), Nbp2-Ste20 (PDB accession

2LCS), and Bem1-2-Ste20 (PDB accession 2RQW) were all previously measured NMR structure ensembles, composed of 20 individual structures. All 20 structures were used for determining the contacts, although only the designated representative structure was used for visualization. The structures were imported into ChimeraX 1.8 (Pettersen *et al.* 2021), and the ChimeraX “Contacts” tool was used to detect all pairs of atoms from different proteins with a Van der Waals radius overlap ≥ -0.40 Å. This was done for all atoms in the motif-bearing protein preceding position -3 in the standard motif numbering. So, in Pbs2, this was done for all atoms in positions 71 to 92 (SNQSEQDKGSSQSPKHIQQIVN), for Ark1 this was done for positions -10 to -4 (KPKLHSP), and for Ste20 this was done for positions -14 to -4 (SSSANGKFIPS). To identify equivalent positions in the four SH3 domains which form contacts, the structures of the 4 domains were aligned using the MUSTANG tool (Konagurthu *et al.* 2006), and a standard numbering was determined based on this alignment (Table S9 in File S2). The contacts between atoms were filtered to simply indicate all contacts between residues, and are summarized in Table S10 in File S2, using the aforementioned standard numbering. The minimum distance between residues or proteins was calculated in a custom R script, using the bio3d package (Grant *et al.* 2006).

### Sequence and structural alignments

The protein sequences of 24 yeast SH3-containing proteins were obtained from the alliancemine server (Bult and Sternberg 2023). The sequence of the SH3 domains plus 25 residues on each side was extracted from the complete protein sequences using the SH3 location information from Interpro (Paysan-Lafosse *et al.* 2023), and a custom R script. When an SH3 domain was within less than 25 residues of the N- or C-terminus of a protein, the sequence up to the terminus was used. The sequences of these SH3 domains and adjoining regions were aligned using mafft 7.526, using the L-INS_i (local alignment) algorithm with the command line : "/usr/bin/mafft" --localpair --maxiterate 16 --inputorder “<input_file>” > “<output_file>” (Katoh and Standley 2013). For the structural alignment, we used previously predicted structures from the AlphaFold Protein Structure Database (Varadi *et al.* 2022). Using the R packages AlphaMissenseR (Nguyen *et al.* 2025) and bio3d (Grant *et al.* 2006), we downloaded the predicted structures of the 24 same yeast SH3-containing proteins from the AlphaFold Protein Structure Database, and trimmed the pdb files to only contains residues composing the SH3 domains as well as 25 residues on each side of the domain. To align the predicted sequences, we used the MUSTANG 3.2.4 tool, with the command line ./bin/mustang-3.2.3 -f ./path_to_files.txt -F fasta -o SH3_flanking -s off (Konagurthu *et al.* 2006). The ./path_to_files.txt file contains the paths of all trimmed pdb files. The sequence and structure alignments in Figure S15 (see File S1) were visualised using the R package ggMSA (Zhou *et al.* 2022).

### Analysis of DHFR-PCA on solid media pictures

The ImageMagick command line tool (ImageMagick Studio LLC 2023) was used to prepare the colony array pictures by cropping the images of the selection plates as well as converting them to grayscale and inverting the colors. The following commands were used: convert -colorspace LinearGray, convert -crop 4200X2800+530+320 and convert -negate. Colony sizes were then quantified using the Python package Pyphe (Kamrad *et al.* 2020), using the command pyphe-quantify timecourse --grid auto_1536 --t 1 --d 3 --s 0.05. These colony sizes were then analysed using a custom script, as detailed in the main text.

Large Language Model use for manuscript editing

The original manuscript draft was entirely written by the authors. Google's large language model Gemini Flash 2.5 was used after manuscript drafting to obtain feedback on grammar and syntax. This was done only to increase the clarity and conciseness of the text. No scientific conclusions were determined by the LLM, and no sources were obtained from the LLM. Suggestions from the model were selectively applied, based on the authors' judgment.
